# Fluorescent Ubiquitination-based Trbl Turnover Indicator ( FUTTI): A tool to detect activity of the Tribbles pseudokinase in vivo

**DOI:** 10.64898/2026.09.25.753906

**Authors:** C. Nauman, S. Shayestehpour, S O’Connor, Z. Fischer, L.L Dobens

## Abstract

A fluorescent sensor detects protein turnover mediated by the Trbl pseudokinase in transgenic flies Abstract (168 words) Targeted protein turnover of the transcription factor C/EBP is enhanced by the Tribbles (Trib) pseudokinase family of adaptor proteins, which bind to a conserved degron motif in C/EBP substrate and recruit a cognate COP1 E3 ligase to ubiquitinate C/EBP for degradation. We identified a conserved degron in the Drosophila C/EBP ortholog Slbo and fused this motif to a fluorescent protein (GFP or RFP) under the control of a constitutive promoter. Transgenic flies expressing this fluorescent protein (FP)::degron transgene, which we call the Fluorescent Ubiquitination-based Trbl Turnover Indicator (or FUTTI) sensor, showed reduced levels of fluorescence in response to increased levels of Drosophila Tribbles (Trbl) and conversely, increased levels when Trbl activity was reduced. Mutations in the Trbl conserved pseudokinase substrate binding pocket and in a divergent PEST-like motif in the C-terminal tail region disrupted Trbl activity in the FUTTI assay. These data indicate that FUTTI will be a valuable tool to identify sites of Trbl activity in vivo and the critical features of the protein required for function.

## Introduction

Focused protein degradation is enhanced by adaptor molecules which bridge substrates to cognate E3 ligases to mediate targeted proteasomal turnover (Zheng and Shabek, 2017; Zhang et al., 2025b; Larsen et al., 2026; Wang and Zheng, 2026). The Tribbles (Trib) family of adaptor proteins this mediate selective protein turnover in all metazoa, with members including the Trib isoforms 1, 2 and 3 in mammals, the NIPI-1 protein in C. elegans and the Tribbles (Trbl) protein in Drosophila (Eyers et al., 2017; McMillan et al., 2021b; Arif et al., 2023; Peng and Wang, 2025; Zhang et al., 2025a). Tribs are related to the calcium/calmodulin-dependent protein kinase sub-family of serine/threonine kinases and share the archetypal DLK catalytic loop motif flanked by (1) an N-terminal lobe that in some mammalian isoforms contains PEST domains, thought to be important for their turnover (Wang et al., 2013; Soubeyrand et al., 2016) and (2) a C-terminal tail, which harbors a MEK1 binding and a COP1 binding motifs, which mediate interactions with MAP kinase kinases and cognate E3 ligases, respectively (Eyers et al., 2017). The crystal structure of Trib1 revealed a protein structurally similar to catalytically active kinases, with an N-lobe connected to a C-lobe via a central cleft region that, unlike the cleft region in bona fide kinases, lacks motifs responsible for both binding ATP and catalytic phosphoryl transfer, and hence the family of proteins have been classified as pseudokinases (Murphy et al., 2015) (Eyers et al., 2017).

In Drosophila, Tribbles was identified in an overexpression screen for factors that influence cell migration in the ovary and was found to act by binding and directing the turnover of the fly version of the C/EBP protein Slbo (encoded by the *slbo* gene; Rørth et al., 2000; Mata et al., 2000; Seher and Leptin, 2000; Großhans and Wieschaus, 2000). This interaction between Tribbles and C/EBP occurs in all animals during normal development and disruptions have been linked to solid tumors and leukemias. In C. elegans, the Trib ortholog NIPI-3 binds and degrades C/EBP in intestinal hypodermis cells to control inflammation and infection resistance in the gut (Kim et al., 2016; McEwan et al., 2016; Wu et al., 2021), and during mouse and human hematopoiesis, Trib1 and Trib2 block neutrophil differentiation in part by facilitating the polyubiquitination of lysine 48 of C/EBPα by the COP1 E3 ligase leading to proteasomal turnover (Keeshan et al., 2006; Bezy et al., 2007; Naiki et al., 2007; Du and Ding, 2009; Keeshan et al., 2010; Xu et al., 2014; Bauer et al., 2015; Richmond and Keeshan, 2020). The pathogenesis of Acute Myeloid Leukemia (AML), a fast-progressing cancer of the blood and bone marrow characterized by the rapid overproduction of immature myeloid cells, is associated with aberrantly high levels of Trib1/2, which are thought to degrade C/EBP and prevent normal blood cell maturation (Keeshan et al., 2006; Jin et al., 2007; Dedhia et al., 2010; Liang et al., 2013; Yoshida et al., 2013; Nakamura, 2015; O’Connor et al., 2016; O’Connor et al., 2018; Sunami et al., 2024; Velasco and Lorente, 2026). The effects of Trib1-induced leukemogenesis in transgenic mice can be reversed by co-expression of ligase activity-deficient COP1 mutant, highlighting how interactions between degrons and E3 ligases bridged by Tribs mediates the pathophysiology of the disease (Keeshan et al., 2010; Yoshida et al., 2013; Murphy et al., 2015; Singh et al., 2024).

Evaluation of changes in Trib1 structure bound or unbound to the substrate C/EBP degradation motif – or degron – has revealed that in its free state, the C-terminal tail of Trib1 is sequestered via an intramolecular binding site in the N-lobe of the protein while binding of the C/EBP degron to a conserved “pseudokinase substrate-like” pocket in Trib1 elicits a conformational change resulting in the release of the C-terminal tail, which recruits a cognate COP1 E3 ligase leading to ubiquitination and rapid turnover of the C/EBP substrate (Murphy et al., 2015; Jamieson et al., 2018; McMillan et al., 2021a). In both HeLa cells and transgenic mice, degrons have been fused to fluorescent proteins to detect interactions between adaptor proteins and E3 ubiquitin ligases (Dantuma et al., 2000; Lindsten and Dantuma, 2003). The best example of this is the FUCCI system (fluorescent ubiquitination-based cell cycle indicator; Zielke et al., 2014; Zielke and Edgar, 2015), in which fluorochrome proteins (FP) tagged with a degron from either the Cyclin B or E2F1 proteins are degraded by the ubiquitin E3-ligase APC/C during M-phase or CRL4Cdt2 E3-ligase at the onset of S-phase, respectively, in proliferating cells (Davoli and de Lange, 2012; Santos et al., 2012; Calder et al., 2013; Choi et al., 2013). To develop a similar tool applicable to the Trib adaptor, we aligned the human C/EBPα degron motif identified by the Mace group (Murphy et al., 2015; Jamieson et al., 2018) to the fly Slbo protein, fused this conserved degron to GFP or RFP and expressed it ubiquitously in transgenic flies. Here we show that this FP::degron can detect sites of Trbl activity in tissues and is sensitive to genetic manipulations that increase or decrease Trbl levels. Further, we show that this FUTTI (Fluorescent Ubiquitination-based Trbl Turnover Indicator) transgene can be used for structure-function assays to identify features of the Trbl protein required for substrate turnover.

## Results

### FUTTI is sensitive to increased and decreased Trbl activity in the wing disc

Structural and mutational analysis of the interaction between human Trib1 and C/EBP has revealed the details of a pseudokinase binding site in the Trib1 protein that serves to interact with a proposed C/EBP degron (Jamieson et al., 2018). This degron motif in human C/EBP is highly conserved among the human isoforms C/EBPα, C/EBPβ, and C/EBPδ, and the conservation among these proteins extends to the Drosophila C/EBP ortholog Slbo, encoded by the *slow border cells* gene (Fig. 1A, Montell et al., 1992; Rorth et al., 2000). We considered the Alphafold structure of Drosophila Slbo to determine that the fly degron motif from residues 118 to 227 was located in an unstructured domain, independent of the C-terminal bZIP DNA binding domain (Fig. 1B; Jumper et al., 2021). We used Alphafold2 to test the in silico interaction between this putative degron region 118-227 and the central domain of Trbl (T59-Y406) and observed specific interactions with the proposed pseudokinase substrate pocket of fly Trbl (Fig. 1C; Jamieson et al., 2018). To determine if this degron could function independently of Slbo in vivo, we fused this domain to either GFP or RFP and placed these sequences under the control of the constitutive ubiquitin promoter in the transgene vectors pUGW or pURW, respectively (Fig. 1D; pUWG and pUWR; see Material and Methods). Verified vectors (FUTTI-GFP and FUTTI-RFP, respectively) were used to construct first and third chromosome transgenic stocks with targeted P-element recombination methods.

**Figure 1.**
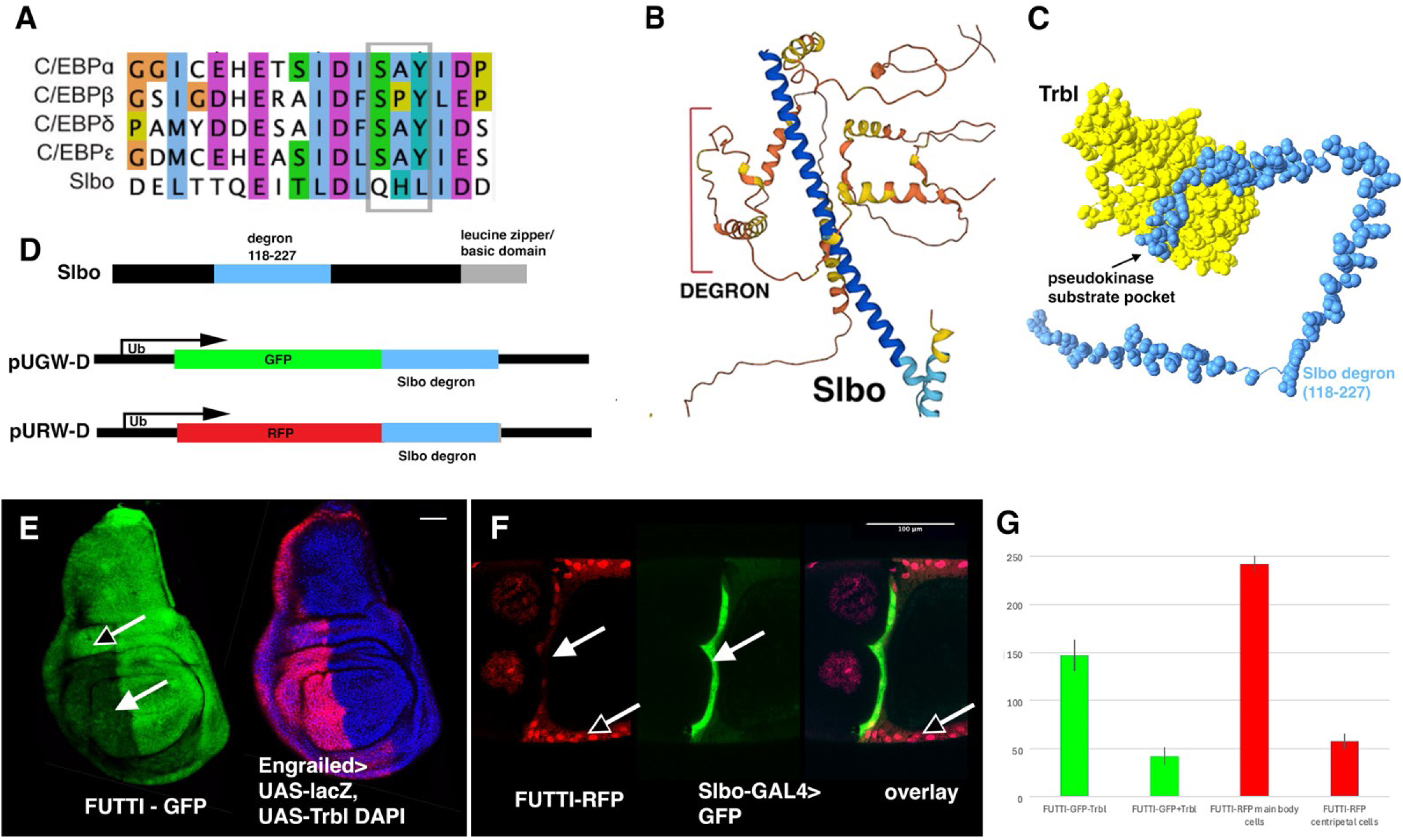
FUTTI System: A Tool for In Vivo Detection of Trbl Activity. (A) CLUSTAL alignment of Trib degron from human C/EBP isoforms and the Drosophila C/EBP ortholog, Slbo (Sievers et al., 2011). Conserved residues are indicated by color and gray box shows mammalian phosphorylation site that is divergent from Slbo. (B) Alphafold structure of Drosophila Slbo protein shows location of degron from 1A. (in bracket) relative to the C-terminal leucine zipper alpha helix (blue ribbon). (C) Alphafold2 interaction between central domain of Trbl (58-206) and the Slbo degron (Slbo_118-227_) reveals a model where the degron (blue) binds the conserved pseudokinase substrate pocket of Trbl (yellow). (D) Construction of FUTTI-GFP and FUTTI-RFP. Fragment (118-227) that contains the conserved predicted degron with flanking amino acids and was inserted into the constructs pUGW and pURW to produce the constructs pUGW-D (degron) and pURW-D, respectively. Transgenic FUTTI strains include this construct with Ubiquitin promoter (Ub) followed by a leader, Shine-Delgarno sequence and GFP::Slbo_118-227_ or RFP::Slbo_118-227_.(E) Wing disks expressing UAS-Trbl in the posterior compartment under control of the Engrailed-GAL4 driver (en2.4-GAL4) marked by UAS-lacZ revealed reduced accumulation of Ub-GFP::Slbo_118-227_ strongly in the wing blade (arrow) and weakly in the dorsal hinge region (empty arrowhead). Scale bar is 100um in this panel and all others. (F) Ub-RFP::Slbo_118-227_ levels are strongly reduced in the centripetally migrating follicle cells (arrow, left) expressing Slbo-GAL4,UAS-GFP (arrow, right). (G) Mean intensity for FUTTI-GFP +/- Trbl expression in the posterior/anterior wing blade tissue and FUTTI-RFP mean intensity in mainbody FC/centripetal FC nuclei expressed in in pixels/area.

To validate these FUTTI system constructs, we first examined the effect of Trbl misexpression on FUTTI-GFP in the posterior compartment of the wing, a tissue sensitive to increased and decreased Trbl levels (Masoner et al., 2013). To do this, we built a stock bearing both the Engrailed promoter fused to GAL4 ({*P*(en2.4-GAL4)}*e16E*; hereafter Engrailed-GAL4 or En-GAL4) and UAS-lacZ and crossed it to a stock bearing FUTTI-GFP and UAS-Trbl (Masoner et al., 2013). As shown in Fig. 1E, Trbl misexpression in the posterior wing blade revealed by UAS-lacZ expression (detected using antisera) reduced FUTTI-GFP in the wing blade and ventral hinge region (solid arrow, Fig. 1E) with FUTTI-GFP intensity reduced in these cells 3.5-fold when expressed in pixels/area (Fig. 1G). We note that in some dorsal hinge and thorax cells, Trbl misexpression resulted in higher levels of GFP expression (e.g. empty arrow in Fig. 1E), suggesting that turnover of the FUTTI-GFP substrate by Trbl is tissue-specific. We evaluated FUTTI-GFP and FUTTI-RFP in the ovary where Trbl regulates Slbo levels in two migrating follicle cell groups: the centripetal cells and the border cells. FUTTI-RFP (Fig. 1F) levels are reduced in the migrating centripetal follicle cells (arrow, 1F) compared to the main body follicle cells (empty arrow, 1F) where FUTTI-RFP remains at high levels. When considering RFP intensity in pixels/area, the reduction is 13.5-fold, (Fig. 1G), suggesting Trbl activity is strong in these migrating cells. Similar results were seen with FUTTI-GFP (data not shown). Together these data support the notion that turnover of GFP or RFP reporters fused to the Slbo degron is sensitive to both endogenous Trbl activity and ectopic increases in Trbl activity.

To further evaluate the effect of changes in Trbl activity on FUTTI-GFP, we constructed a stock bearing Engrailed-GAL4, UAS-RFP and FUTTI-GFP and crossed this test stock to UAS-regulated transgenes designed to increase or decrease Trbl activity (Fig. 2). FUTTI-GFP expression in the wing disc is ubiquitous (Fig. 2A) and Engrailed-GAL4 expression of Trbl as detected by UAS-RFP reduced FUTTI-GFP levels in the posterior compartment (Fig. 2B, arrow). By contrast, when we reduced Trbl activity in posterior cells using Engrailed-GAL4 expression of either Trbl RNAi (Fig. 2C) or a dominant negative version of Trbl that forms unproductive Trbl multimers (UAS-Flag-Trbl^286E/G^ ; Fig. 2D; Fischer et al., 2025), we observed increased FUTTI-GFP accumulation (Figs 2C and D, arrows).

**Figure 2.**
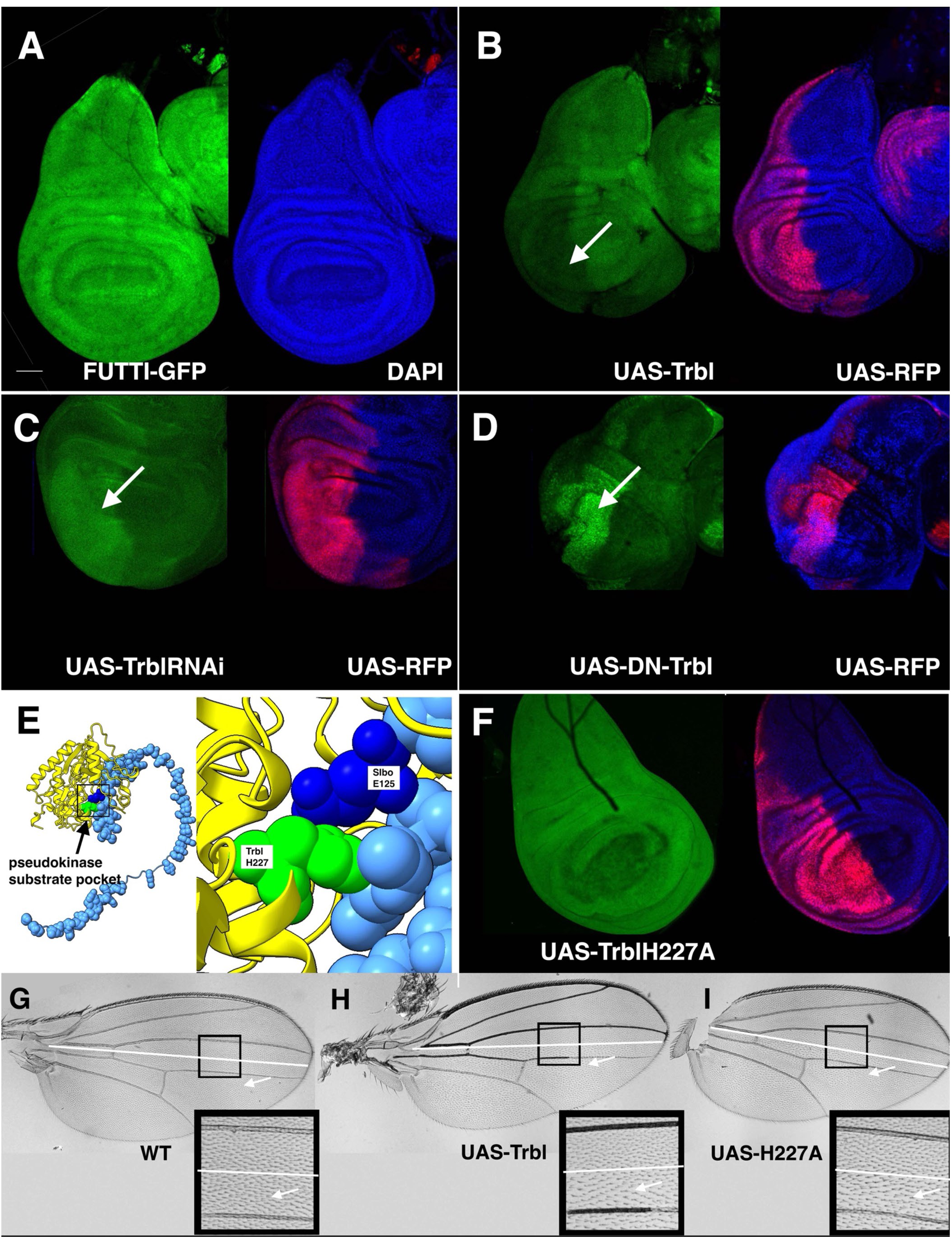
FUTTI-GFP is Sensitive to Increased and Decreased Trbl Levels. (A) Wing disc cells expressing *Ub*-GFP::Slbo_118–227_ from the Ubiquitin promoter showed uniform GFP accumulation throughout the wing disk. DAPI in blue here and in all panels. (B) Wing disk expressing UAS-Trbl under the direction of the Engrailed-GAL4 driver expressing UAS-RFP (red, here and in panels B,C,D and F) revealed reduced Ub-GFP::Slbo_118-227_ levels (arrow) in the posterior compartment, strongly in the wing blade and reduced in the wing hinge regions. (C) Engrailed-GAL4>UAS-Trbl^22114|GD^ RNAi revealed increased Ub-GFP::Slbo_118-227_ levels (arrow) in the posterior compartment. (D) Engrailed-GAL4>UAS-DN-Trbl (Trbl^E286G^, a dominant negative version of Trbl) revealed increased Ub-GFP::Slbo_118-227_ levels (arrow) in the posterior compartment. (E) Alphafold2 interaction between central domain of Trbl (58-206, yellow ribbon) with the Slbo degron (Slbo_118-227_, space filling in light blue) reveals binding at pseudokinase pocket (arrow). Inset shows predicted proximity of TrblH227 (green) with SlboE125, in the degron (dark blue). (F) Engrailed-GAL4>UAS-Trbl^H227A^ mutation revealed no reduction in Ub-GFP::Slbo_118-227_ levels in the posterior compartment. (G) Engrailed-GAL4>expression of UAS-lacZ showed WT distribution of trichomes in the posterior wing (inset, arrow). The boundary between anterior and posterior compartments are indicated. (H) Engrailed-GAL4>expression of UAS-Trbl showed reduced density of trichomes in the posterior wing (inset, arrow). (I) Engrailed-GAL4>expression of UAS-Trbl^H227A^ showed reduced density of trichomes in the posterior wing.

Finally, to test the specificity of Trbl for the FUTTI-GFP degron, we mutagenized the pseudokinase substrate-like pocket, specifically a residue H227 that corresponds to a conserved histidine in human Trib1 (H168) shown to be required for interactions with C/EBP (Fig. 2E; Jamieson et al., 2018). Engrailed-GAL4 expression of an alanine replacement in H227 (UAS-Trbl^H227A^) resulted in no reduction in FUTTI-GFP in the posterior compartment of the wing (Fig. 2F). This result confirms the critical role of the pseudokinase substrate-like pocket for FUTTI substrate turnover. Unexpectedly Engrailed-GAL4> expression of the pocket mutation Trbl^H227A^ resulted a reduced density of trichomes in mature wings similar to expression of wild type (WT) Trbl with an intact pocket (cf. Fig. 2G and 2H and insets). This suggests that while the pseudokinase substrate-like pocket mutation Trbl^H227A^ has lost its ability to interact with Slbo in the FUTTI assay, Trbl^H227A^ retains its ability to block String/Twine (Cdc42 phosphatase)-mediated cell division, reducing the density of trichomes.

### Mammalian Trib isoforms do not direct FUTTI turnover

In mammals, Trib isoforms direct C/EBP turnover via the conserved COP1 E3 ubiquitin ligase, a member of an ancient family of E3 ubiquitin ligases that binds Trib isoforms via a WD40 domain and E2 ubiquitin-conjugating enzyme via a conserved N-terminal RING domain to catalyze exchange of a ubiquitin moiety to a bound substrate (reviewed in Yi and Deng, 2005). As shown in Fig. 3, misexpression of mouse Trib isoforms Trib1, (Fig. 3A) Trib2 (Fig. 3B) and Trib3 (Fig. 3C) in the posterior compartment did not reduce FUTTI-GFP expression. This was a surprise given the conserved nature of the degron, which would lead us to predict that mouse Trib isoforms might interact with and direct turnover of the Slbo degron.

**Figure 3.**
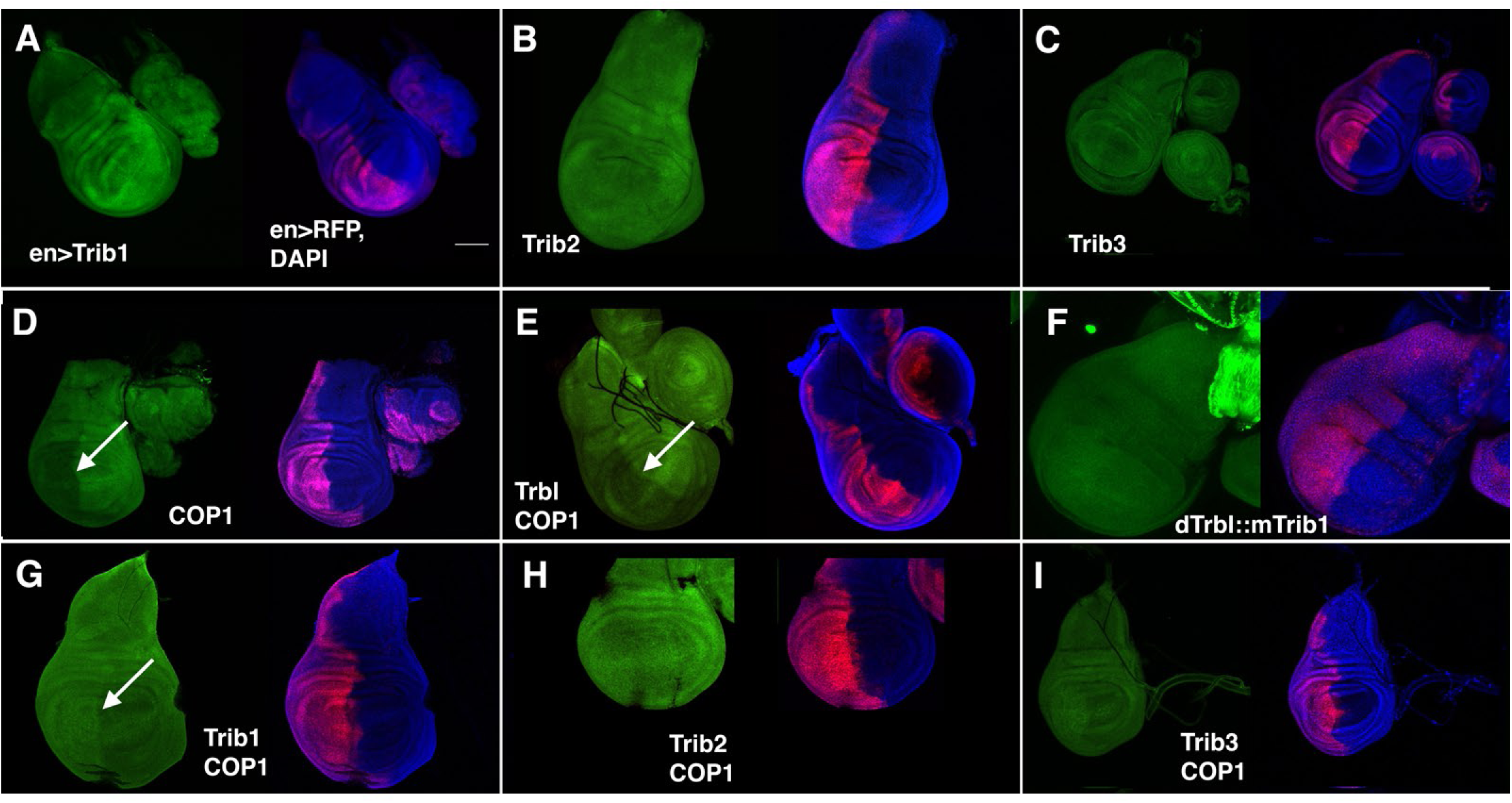
FUTTI-GFP is Insensitive to Expression of Mammalian Trib Isoforms. (A) Wing disk expressing UAS-Trib1, the mouse homolog of Tribbles, under the direction of the Engrailed-GAL4 driver expressing UAS-RFP (red here and in all panels) revealed no change in Ub-GFP::Slbo_118-227_ levels in the posterior compartment. DAPI is blue in this and all panels. (B) Engrailed-GAL4>UAS-Trib2, the mouse homolog of Tribbles, revealed no change in Ub-GFP::Slbo_118-227_ levels in the posterior compartment. (C) Engrailed-GAL4>UAS-Trib3, the mouse homolog of Tribbles, revealed no change in Ub-GFP::Slbo_118-227_ levels in the posterior compartment. (D) Engrailed-GAL4> expression of UAS-COP1, the mouse homolog of COP1 E3 ligase revealed reduced Ub-GFP::Slbo_118-227_ levels (arrow) in the posterior compartment, strongly in the wing blade and reduced in the wing hinge regions. (E) Engrailed-GAL4> co-expression of UAS-COP1 and UAS-Trbl showed reduced Ub-GFP::Slbo_118-227_ levels (arrow) in the posterior compartment. (F) Engrailed-GAL4> expression of UAS-Trbl^I391Trib1^, a replacement of the Trbl C-terminal tail with the mouse Trib1 C-terminal tail, showed slight increase in Ub-GFP::Slbo_118-227_ levels (arrow) in the posterior compartment (G) Engrailed-GAL4> co-expression of UAS-COP1 and UAS-Trib1 showed slightly increased Ub-GFP::Slbo_118-227_ levels (arrow) in the posterior compartment. (H) Engrailed-GAL4> co-expression of UAS-COP1 and UAS-Trib2 showed similar Ub-GFP::Slbo_118-227_ levels (arrow) in the posterior compartment compared to the anterior compartment. (I) Engrailed-GAL4> co-expression of UAS-COP1 and UAS-Trib1 showed increased Ub-GFP::Slbo_118-227_ levels (arrow) in the posterior compartment.

We and others noted previously that no COP1 E3 ligase homolog exists in Drosophila (Dobens et al., 2021), so to test if the mammalian COP1 E3 ubiquitin ligase might potentiate mouse Trib isoform function in the FUTTI assay, we inserted the mouse COP1 open reading frame into the pUAS-T vector and generated transgenic flies expressing mouse COP1 under control of the UAS-promoter (materials and methods). Engrailed-GAL4 driving expression of UAS-mCOP1 in the posterior wing compartment reduced FUTTI-GFP substrate levels (Fig. 3D). However, co-expression of COP1 with either Trib1, (Fig. 3G), Trib2 (Fig. 3H) or Trib3 (Fig. 3I) did not further reduce FUTTI-GFP levels, but rather these transgenes expressing mammalian Trib isoforms suppressed COP1-mediated turnover of FUTTI, resulting in higher GFP levels. In contrast, Trbl co-expression with COP1 does not suppress the effects of COP1 expression, resulting in lower levels of GFP (Fig. 3E) and supporting the notion that Trbl does not interact with COP1. These results suggest that mammalian Trib isoforms do interact with mammalian COP1 E3 ligase but do so unproductively in the FUTTI wing assay.

### Structure-function analysis of the Trbl C-terminal tail reveals a motif unique to the Drosophila lineage required for FUTTI turnover

Trib1-COP1 co-crystals reveal that the COP1 WD40 beta-propeller domain has a small hydrophobic surface pocket off-center from the central doughnut hole that binds a valine and proline found in the conserved COP1 binding sites (E/DQxVPE/D) found in the three human Trib isoforms (Uljon et al., 2016). We tested a point mutation Trbl^E436A^ in the proposed COP1 binding site of fly Trbl (‘EGLCP’; Dobens and Bouyain, 2012; Yokoyama and Nakamura, 2011b), and found this mutation retained Trbl function to effectively degrade FUTTI substrate (Fig. 4E). When we replaced the Trbl C-terminal tail with that of mouse Trib1, fusing the Trbl protein from 0-390 in frame with the C-terminal region of mammalian Trib1 (residues 332-372; including the MEK1 and COP1 binding domains), we found that this chimera had no activity in the FUTTI assay (Fig. 3F). We surmise that while Trbl directs FUTTI turnover, it does so via an E3 ubiquitin ligase distinct from COP1.

**Figure 4.**
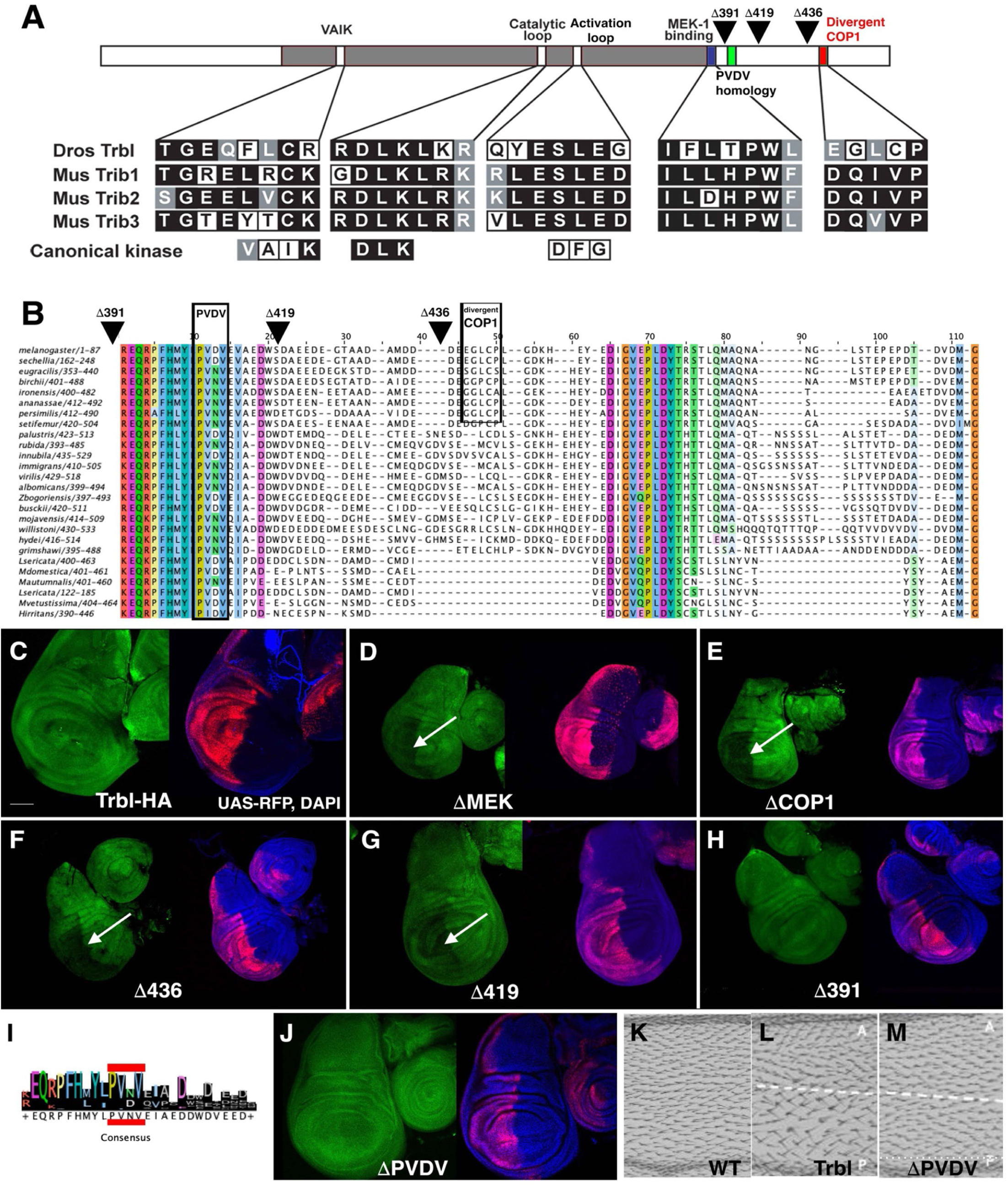
Structure-function Analysis of Trbl using FUTTI-GFP Identifies a Divergent Domain required for Substrate Turnover. (A) Schematic of Trbl protein with N-lobe (white), central pseudokinase domain (gray) and C-terminal tail (white) with conserved domains highlighted, including the divergent VAIK domain, the divergent activation loop domain and conserved catalytic domain. The C-tail retains a conserved MEK binding site and divergent COP1 binding site. The location of a conserved domain (PVDV) is indicated. Location of deletion mutations D391, D419 and D436 are indicated. (B) COBALT (Constraint-Based Alignment Tool) multiple sequence alignment reveals conserved domains in the Drosophila C-terminal tail with related Diptera. Location of deletion mutations Δ391, Δ419 and Δ436 are indicated. (C) Wing disk expressing UAS-Trbl-HA, a version of Trbl that carries a C-terminal HA tag, under the direction of the Engrailed-GAL4 driver co-expressing UAS-RFP revealed no reduced Ub-GFP::Slbo_118-227_ levels (arrow) in the posterior compartment. (D) Engrailed-GAL4>expression of UAS-Trbl (ΔMEK) showed reduced Ub-GFP::Slbo_118-227_ levels (arrow) in the posterior compartment. (E) Engrailed-GAL4>expression of and UAS-Trbl (ΔCOP1) showed reduced Ub-GFP::Slbo_118-227_ levels (arrow) in the posterior compartment. (F) Engrailed-GAL4>expression of and UAS-Trbl (Δ436) showed reduced Ub-GFP::Slbo_118-227_ levels (arrow) in the posterior compartment. (G) Engrailed-GAL4>expression of and UAS-Trbl (Δ419) showed somewhat reduced Ub-GFP::Slbo_118-227_ levels (arrow) in the posterior compartment. (H) Engrailed-GAL4>expression of and UAS-Trbl^ΔI391^ (Δ391) showed no reduced Ub-GFP::Slbo_118-227_ levels (arrow) in the posterior compartment. (I) Consensus residues in proximal tail region PVDV motif common to related Diptera in panel 4B. (J) Engrailed-GAL4>expression of and Trbl^PVDV408AGKG^ showed no reduced Ub-GFP::Slbo_118-227_ levels (arrow) in the posterior compartment. (K) Engrailed-GAL4>expression of UAS-lacZ showed WT distribution of trichomes in the posterior wing. (L) Engrailed-GAL4>expression of UAS-Trbl showed reduced density of trichomes in the posterior wing. The boundary between anterior and posterior compartments are indicated by white dotted line. (M) Engrailed-GAL4>expression of UAS-Trbl^PVDV408AGKG^ showed no reduced density of trichomes in the posterior wing.

The failure of the putative COP1 binding site mutation Trbl^E436A^ to reduce FUTTI-GFP (Fig. 4E) indicated that the C-terminal tail of Trbl may direct substrate turnover via a novel mechanism. The Drosophila tail has a structure distinct from mammalian isoforms and is more that 40 amino acid residues longer than the Trib3, the longest human Trib protein with a 50-residue tail. Addition of an HA tag to the C-terminal tail resulted in a version of Trbl (Trbl-HA) that fails to direct FUTTI substrate turnover (Fig. 4C), indicating the tail structure loses activity when modified in this way.

To understand the function of the tail, we produced a series of deletion mutations that progressively removed portions of the C-terminal tail of Trbl and tested these in the FUTTI wing assay. Fig. 4A and 4B detail the location of Trbl deletion mutations (Trbl^ΔI391^, Trbl^ΔD419^, Trbl^ΔE436^) that by western blot analysis of larval extracts were confirmed to be expressed in vivo (data not shown). The Trbl C-tail deletion mutation removing residues after residue 436, including both the poorly conserved COP1 binding site (‘divergent COP1’ in 4B) and some sequence motifs conserved among wine flies, retains the ability to direct FUTTI turnover (Fig. 4F). Further deletions to residue 419 also demonstrates FUTTI substrate turnover function (Fig. 4G) and only complete deletion of the C-tail up to the MEK1 binding site at residue 391 loses all FUTTI activity (Fig. 4H). Because a mutation in the MEK1 binding site itself also has FUTTI activity (Trbl^W386A^; Fig. 4D), we conclude that a 28 amino acid region (391-419) distal to the MEK1 binding site is required for full FUTTI-GFP turnover.

In a quest to broaden our understanding of the evolutionary context of *D. melanogaster* Tribbles C-tail, we collected arthropod Trbl sequences from the NCBI database and generated a parsed list trimmed down to just the C-terminal tail (Fig. 4B). Based on this alignment, we determined that the putative COP1 binding site was not conserved at all (Fig. 4B), however we detected two conserved motifs common to the wine fly lineage (Fig. 4B). One of these domains contains a conserved PVDV motif (Fig. 4I) from 408-411 that we replace with the residues AGKG in the context of a full length Trbl sequence. Engrailed-GAL4 expression of UAS-Trbl^PVDV408AGKG^ in the posterior wing resulted in no reduction in FUTTI-GFP levels (Fig. 4J), consistent with a requirement of this conserved PVDV motif for turnover. We have shown previously that WT Trbl expressed in the posterior wing leads to reduced trichome density (cf. Fig. 4L and M), and Engrailed-GAL4 expression of UAS-Trbl^PVDV408AGKG^ mutation abrogated this phenotype, indicating that the PVDV motif in the C-tail is critical for recruiting the proteasome for both Slbo and String/Twine turnover.

## DISCUSSION

### FUTTI-GFP and -RFP detect Trbl activity in vivo

The diverse roles for Tribbles family members in cell proliferation, differentiation, and tissue homeostasis relies on two conserved features: (1) a pseudokinase substrate-like pocket in the central domain, which is predicted to bind degron motifs in substrates, and (2) a COP1 binding site at the C-terminal region, that binds E3 ubiquitin ligases leading to ubiquitination and degradation of the substrate (Keeshan et al., 2010; Yokoyama and Nakamura, 2011a). The interaction between Trib1 and the internal degron of C/EBP elicits a conformational change to expose an E3 ubiquitin ligase binding site in the C-terminal tail, recruiting E3 ubiquitin ligase to the substrate-bound compled, leading to E2 enzymatic ubiquitination and proteasomal degradation of the target protein. We have exploited the deep conservation of the human C/EBP degron with the Drosophila C/EBP ortholog Slbo (Slow Border Cells) and fused the predicted fly Slbo degron to the fluorescent proteins GFP and RFP in a transgenic construct These tools, which we call **F**luorescent **U**biquitination-based **T**rbl **T**urnover **I**ndicators (or **FUTTI**), show that FP::degron levels vary inversely with endogenous Trbl activity in ovarian tissue and are sensitive to increased and decreased Trbl levels in the wing model.

### Structure-function analysis of Trbl evaluated by FUTTI activity

The ability of Trbl to efficiently direct FUTTI substrate turnover led us to use this tool to evaluate the functional features of the Trbl protein required for activity. The pseudokinase binding pocket in Trib1 has been shown by extensive mutagenesis to be essential for C/EBP binding and by comparing the Alphafold structures of Trib1 to fly Trbl, we identified a conserved histidine residue in the binding pocket shared by both proteins (H168 in human Trib1 and H227 in fly Trbl). Expression of the Trbl^H227A^ resulted in no significant FUTTI substrate turnover, defining better the interface between Trbl and the Slbo degron.

Complementarily, we mutagenized the C-terminal portion of Trbl to identify residues required for E3 ubiquitin ligase binding. In diverse metazoans Trib isoforms display a COP1 binding consensus sequence E/DQxVPE/D in the C-terminal tail that is important for high affinity binding of the conserved E3 ubiquitin ligase COP1 (Uljon et al., 2016; Kung and Jura, 2019). Experimental data validates the importance of this motif for mammalian Trib proteins: a mutation in human Trib1 that disrupts the COP1 binding motif is unable to direct C/EBP turnover (Qi et al., 2006; Keeshan et al., 2010; Yokoyama and Nakamura, 2011a). We previously flagged a possible COP1 binding site in the C-terminal tail of Trbl (Dobens and Bouyain, 2012), but mutagenesis of this motif (Trbl^E436A^) did not affect the protein’s ability to direct FUTTI-GFP turnover. While this result was unexpected, upon closer examination we concluded that Drosophilids (and eleven other Dipterans of the suborder Schizophora) apparently lack a clear COP1 consensus sequence, and consistent with this, complete deletion of this motif in the deletion mutant Trbl^ΔD419^ resulted in a version of Trbl that nevertheless had strong FUTTI activity.

We and others have noted previously that Drosophila apparently has no ortholog of the COP1 E3 ubiquitin ligase among its 250 catalogued E3 ubiquitin ligases (Dobens et al., 2021). Consistent with this, (1) a chimeric Trbl-Trib1 protein, where the entirety of Trbl C-terminal tail is replaced with that of mouse Trib1, was unable to perform Trbl-directed degradation, and (2) none of the mouse Trib isoforms that do rely on COP1 ligase for function direct FUTTI turnover in the Drosophila wing model. Ectopic expression of mouse COP1 alone did lead to reduced FUTTI GFP, indicating that FUTTI is sensitive to increased levels of this E3 ubiquitin ligase. However, when we co-expressed mouse COP1 with Trib isoforms (1, 2 and 3), we observed no potentiation of the effect of mCOP1 on FUTTI turnover, but rather suppression leading to increased levels of GFP. These data indicate that mouse Tribs and COP1 do interact in Drosophila wing cells, but the failure to direct FUTTI turnover suggests Tribs might be selective for mouse-specific degrons.

To understand the functional features particular to the fly C-terminal tail, we performed a CLUSTAL evaluation of tails distal to the conserved MEK1 site in related Drosophilids to identify a novel motif, which we have called the PVDV motif. This motif is found in the *Drosophilidae* and several non-*Drosophilidae* Dipterans of the suborder Schizophora but absent in the Arthropoda and all other metazoan tails that include the mammals. When the PVDV motif is deleted in the Trbl^ΔI391^ mutant or subject to site-specific mutagenesis in the Trbl^PVDV408AGKG^ mutant, FUTTI substrate turnover was lost. It is worth noting that while FUTTI turnover similarly fails to occur by mutations in the pseudokinase pocket (Trbl^H227A^) and the C-terminal tail (Trbl^PVDV408AGKG^), the pocket mutation retains activity in the wing trichome assay while the tail mutation does not. This suggests that the C-terminal tail that binds E3 ligase is required generally to mediate proteasomal degradation of all substrates, while the pseudokinase substrate-like pocket is selective for the C/EBP target.

The functional requirement for the PVDV motif in lieu of a typical COP1 binding site suggests that Trbl has adopted a novel mechanism for recruiting the ubiquitination machinery to direct substrate turnover. An evaluation of the Trbl protein for PEST domains using the EMBOSS epestfind online tool, which generates a PEST score by combining a sequences P, E, and S/T enrichment, as a mass percentage, and hydrophobicity index (Rogers et al., 1986), identified a strong PEST motif (locus 404 to 444) located in the region spanning the conserved PVDV motif. The PEST motif was first identified in proteins that undergo rapid turnover and are marked by a hydrophilic stretch flanked by positive amino acids with at least one proline (P), one glutamic acid (E), and one serine (S) or threonine (T) (reviewed in Rechsteiner and Rogers, 1996; Sarfraz et al., 2021). The vast majority of Drosophilid Trib proteins have this PEST binding motif encompassing the PVDV region, with 93% of members having PEST scores above zero and 60% having scores above five, a typical threshold for valid PEST domains (data not shown). While the precise mechanism of PEST domain action has remained unclear, PEST domains required for turnover of HSF2 (Heat shock factor 2) were found to bind directly the Cullin 3 (Xing et al., 2010), a core component of the Cullin complex that connects adaptor proteins to E2 proteins charged with ubiquitin moieties to tag these for degradation (reviewed in Nguyen et al., 2017; Nguyen et al., 2017). It is possible that this putative PEST domain in fly Trbl C-tail interacts directly with a Cul3-like component of the proteasome complex, obviating the need for a COP1 ligase homolog to direct turnover and placing fly Trbl in a novel class of E3 ligase-independent proteasomal adaptors such as Stuxnet (Midnolin; Du et al., 2016; Zhu et al., 2026).

## Materials and Methods

### Constructs

Site directed mutant versions of Trbl were generated from pUAST-attB-FLAG-Trbl (Masoner *et al*., 2013) using the QuikChange II XL Site-Directed Mutagenesis Kit (Stratagene) to generate Trbl^E436A^ and Trbl^W396A^ or the Q5 Site-Directed Mutagenesis Kit (NEB) to generated Trbl^Δ419^, Trbl^PVDV408AGKG^, Trbl^ΔE436^, and Trbl^ΔI391^ (see Table 2.3 for primers used). Trbl^I391Trib1^ chimeric gene was GBLOCK (GeneScript) designed with restriction sites to EcoRI and KpnI and cloned into the pUAST-attB vector. Site-directed mutagenesis (lower case letters in primer sets) was confirmed by Sanger Sequencing performed by MU-DNA Core.

UAS-mCOP1 was synthesized by G-BLOCK (Twist Bioscience) designed by the mouse ORF with Drosophila codon preference flanked by Gateway cloning sites and inserted into the pUAS-T Gateway compatible construct.

Embryos were injected as a fee-for-service (Trbl^E436A^ and Trbl^W396A^ by Genetic Services, Inc., Cambridge, MA; Trbl^ΔE436^ and Trbl^ΔI391^ by Rainbow Transgenic Flies, Inc., Camarillo, CA; and Trbl^I391Trib1^, Trbl^PVDV408AGKG^, and Trbl^ΔD419^ by GenetiVision Corporation, Houston, TX) into the landing site attP2 (/68A4 on 3L; Groth et al., 2004).

### Drosophila lines used

1. *w[*]; P*[+mC]=GAL4-slbo.2.6}*1206 P*[+mC]=UAS-GFP.S65T}*T2/CyO* (gift from Dr. Pernille Rorth)
2. *y[1] w[*]; P*[+mW.hs]=en2.4-GAL4}*e22c/SM5*
3. *Sco/Cyo; P*[+mc]=UAS-FLAG-TrblΔE436stop}*3L-68A* (this publication)
4. *Sco/Cyo; P*[+mc]=UAS-FLAG-TrblE436A}*3L-68A* (Masoner et al., 2013)
5. *Sco/Cyo; P*[+mc]=UAS-FLAG-TrblΔI391stop}*3L-68A* (this publication)
6. *Sco/Cyo; P*[+mc]=UAS-FLAG-TrblW396A}*3L-68A* (Masoner et al., 2013)
7. *Sco/Cyo; P*[+mc]=UAS-FLAG-TrblΔD419stop}*3L-68A* (this publication)
8. *Sco/Cyo; P*[+mc]=UAS-FLAG-TrblPVDV408AGKG}*3L-68A* (this publication)
9. *P*[+mc]=UAS-FLAG-Trbl}*3L-68A* (Masoner *et al.,* 2013)
10. *P*[+mc]=UAS-FLAG-Trib1}*3L-68A* (Masoner *et al.,* 2013)
11. *P*[+mc]=UAS-FLAG-Trib2}*3L-68A* (Masoner *et al.,* 2013)
12. *P*[+mc]=UAS-FLAG-Trib3}*3L-68A* (Masoner *et al.,* 2013)
13. *Sco/Cyo; P*[+mc]=UAS-FLAG-TrblΔI391Trib1(murine)}*3L-68A* (this publication)
14. *y1 w*; P(r4-GAL4)3* was a generous gift from Dr. Laura Musselman.

### Immunostaining

Antibodies used to stain tissues: 1:800 rabbit anti HA (Cell Signaling Technologies), 1:1000 rabbit anti-GFP (Sigma), 1:200 mouse anti-b-galactosidase (Developmental Studies Hybridoma Bank, DHSB). Secondary antibodies used were: AlexaFluor 1:200 goat anti-rabbit 594, 1:800 goat anti-mouse 488, and 1:200 goat anti-mouse 594 (Invitrogen). Nuclei were labeled with 1:1000 DAPI (Roche). Wing discs were fixed in a formaldehyde solution (4% paraformaldehyde diluted in PBS) for 30 minutes, washed three times for 5 minutes in NP-40 wash (0.5% IGEPAL, 150 mM NaCl, 50 mM Tris-Cl (pH=7.4). For antisera staining, tissue was blocked in 1mg/mL BSA in H_2_O, 0.1% Triton-X-100, 2% BSA, and 2% NGS in PBS solution) for 2 hours, stained overnight with 1° antibody in PAT solution (0.5% Triton-X-100 and 0.5% BSA diluted in PBS), stained for 2 hours in 2° antibody in PAT solution, washed three more times while adding DAPI. Images were collected on Nikon AXR Resonant Scanning confocal microscope using 10x and 20x objectives and subsequently processed and analyzed with FIJI.

### Phylogenic Tree and WebLogo

*Drosophila* Tribbles protein BLAST was run against NCBI database of Arthropoda (taxid:6656) and Drosophilidae (taxid:7214). The Non-redundant database was used with the following settings: Expect threshold of 10, Word size of 6, Matrix-BLOSUM62, Gap Costs-Existence:11 extension:1, Computational adjustments-Conditional Score matrix adjustment. Sequences were then eliminated from the data set that did not retain important defining features of the Tribbles pseudokinase core. Each sequence was then trimmed to only represent the C-terminal tail as marked by the MEK1 binding site at the N-terminal side. Dash aligned sequences from NCBI were displayed in WebLogo (Crooks, Hon, Chandonia, & Brenner, 2004; Schneider & Stephens, 1990) and edited to represent only residues that aligned to *Drosophila* Tribbles sequence.

### Plasmid construction details

#### UAS-mCOP1

We synthesized a G-Block (below) corresponding to the mouse COP1 open reading frame flanked by a Kozak consensus sequence for efficient translation and a 3’UTR tubulin alpha 1 trailer was used. This was directionally cloned using GATEWAY cloning into pCaSpeR-DEST3 and inserted in attP2 on chr. III.

TCAACATGAGTGGCTCCAGACAAGCGGGTTCCGGATCGGCAGGAACTTCCCCGGGTTCAAGCGCTGCTTCGAGTGTCACGAGTGCTTCGTCCTCGCTGAGCAGTTCCCCCTCACCCCCTAGCGTAGCCGCTAGTGCCGCTACACTCGTTAGCGGTGGTGTTGCGCCAGCGGCAGGATCGGGTGGACTGGGCGGACCTGGTAGACCAGTCCTAGTTGCCGCAGCCGTTTCAGGTAGTGCCTCGGCGGGTGGAGCTGTTAGCGCTGGTCAATCGCGCTTGAGTTGTGCCGCTCGACCATCGGCTGGTGTGGGCGGATCCAGCTCATCACTGGGATCTTCGTCCCGTAAACGCCCGCTACTGGTACCCCTGTGCAATGGTTTGTTGAATAGCTATGAAGATAAGTCGAATGATTTTGTATGCCCAATTTGTTTCGACATGATCGAAGAAGCCTATATGACGAAGTGCGGACATTCCTTCTGTTATAAATGCATCCACCAAAGCCTTGAAGATAACAACCGCTGCCCTAAATGCAACTACGTCGTAGATAACATCGATCACCTATACCCCAACTTTCTGGTAAACGAGCTTATACTGAAGCAAAAGCAGCGCTTCGAAGAGAAACGTTTTAAGCTCGATCATAGCGTATCGTCGACAAACGGACACCGGTGGCAGATCTTCCAGGACCTGCTGGGTACGGACCAGGACAATTTGGACCTCGCTAACGTGAATTTGATGCTGGAGCTGCTGGTTCAGAAGAAGAAGCAGCTTGAGGCCGAGTCCCACGCTGCCCAACTTCAAATATTGATGGAGTTTCTAAAAGTAGCCCGGCGCAACAAACGCGAACAGCTGGAACAAATTCAAAAGGAGCTGTCCGTCCTAGAGGAAGACATCAAGCGTGTCGAGGAGATGAGCGGTCTGTATTCACCAGTGTCGGAAGACAGTACCGTCCCACAGTTCGAGGCCCCCTCACCCAGTCATAGCTCGATAATCGACTCGACCGAGTATTCGCAGCCACCGGGATTTTCAGGCACGTCGCAAACTAAGAAGCAACCCTGGTACAATTCAACCCTCGCGTCTCGTCGCAAAAGGCTTACAGCCCACTTCGAGGATTTGGAACAATGCTACTTCAGCACGCGTATGAGCCGAATAAGCGACGATTCACGAACCGCCTCGCAACTTGACGAGTTCCAGGAGTGTCTGAGCAAATTCACGAGATATAATAGCGTGCGACCACTAGCTACGCTATCGTACGCGTCTGACCTTTACAACGGCAGTTCTATTGTCTCGTCCATAGAATTCGATC

GCGACTGCGATTACTTCGCCATCGCCGGCGTCACGAAGAAAATCAAGGTTTACGAATACGGAACCGTTATTCAAGACGCGGTCGACATCCACTATCCGGAAAACGAGATGACGTGTAACTCTAAGATATCGTGCATTTCATGGAGCTCATATCACAAGAATCTACTGGCTTCATCAGACTACGAGGGAACCGTGATTCTTTGGGACGGCTTTACCGGTCAACGGAGCAAAGTATACCAAGAACACGAGAAGCGTTGCTGGTCGGTGGATTTCAACCTCATGGACCCGAAGCTGCTCGCCAGCGGCTCCGACGACGCCAAGGTCAAACTCTGGAGCACAAACCTGGATAACTCGGTTGCCTCTATCGAAGCTAAAGCAAATGTTTGTTGCGTAAAATTTTCCCCTTCATCACGCTATCACCTCGCCTTTGGTTGCGCGGACCATTGCGTTCATTATTATGATCTGCGGAATACGAAGCAACCCATCATGGTGTTTAAAGGCCATCGCAAGGCTGTTTCCTACGCTAAATTCGTTTCCGGAGAGGAGATAGTGTCGGCTTCGACCGATTCCCAGCTGAAATTGTGGAACGTAGGTAAGCCCTATTGTCTCCGCAGTTTTAAAGGACACATCAACGAGAAGAATTTCGTCGGACTCGCGAGCAACGGTGACTACATCGCTTGCGGCTCAGAAAATAATTCGCTGTATCTATACTACAAGGGCCTGTCGAAGACCCTGCTGACGTTCAAATTCGACACAGTGAAAAGCGTGCTAGATAAGGACCGCAAAGAGGACGATACCAACGAGTTCGTTTCGGCCGTCTGCTGGCGAGCGCTTTCGGATGGCGAATCGAACGTACTCATAGCAGCCAATTCTCAAGGCACCATCAAAGTCCTCGAGCTCGTTTAG

#### FUTTI-GFP

We synthesized a G-Block sequence (below) corresponding to the Slbo open reading frame flanked by GATEWAY cloning site for directional cloning into pUGW and pURW, allowing in-frame fusion with the C-terminus of GFP and RFP, respectively. Following injection P-element transgenic stocks were recovered on the first and third chromosome and strains were inspected for GFP and RFP.

CCTTGTTGCAGAAGCAAATGTTGCAGCAGTACAGTACGCAGACAGATTTGGATGAACTGACGACACAGGAGATTACTTTGGATCTTCAGCACCTCGCAGCCGACCAATTCCGGGATACAGAAACCCTCGGCATATTCTCGGACATGGTGACCTCCCCCGGTGGACTGTCAGCTACGCTACCACCCAGCGGCATGGTGAGTGCCGCCGCGAAAGTGCTTCAGCAGCAAACACTTAGGAATCAGCACGGCTATGGCGGTAGGGGAGGTGGCGGCGGAGCGGGCGGCGCCTTGGCGTATATGCCGCAACCGGTTCACGCAACCTACAACAATTCGAGCGACGAGAATTCCAGCGTTGGCTCCGACAGCTCTACAATTAAGGAAGAACCCATTGACTAA

## Acknowledgments

We thank colleagues for helpful comments on this work. We thank former members of the Dobens lab and students in the BIO312 course for constructing the COP1 expression construct and for helpful comments on earlier versions of the manuscript.

## Data and resource availability

Stocks and constructs will be made available upon request.

